# Omic-Sig: Utilizing Omics Data to Explore and Visualize Kinase-Substrate Interactions

**DOI:** 10.1101/746123

**Authors:** T. Mamie Lih, David J. Clark, Hui Zhang

## Abstract

Protein phosphorylation is one of the most prevalent post-translational modifications, resulting from the activity of protein kinases phosphorylating specific substrates. Multiple cellular processes are regulated via protein phosphorylation, with aberrant signaling driven by dysregulation of phosphorylation events and associating with disease progression (e.g., cancer). Mass spectrometry-based phosphoprotomics approaches can be leveraged for studying alterations of kinase-substrate activity in clinical cohorts. However, the information gained via interrogation of global proteomes and transcriptomes can offer additional insight into the interaction of kinases and their respective substrates. Therefore, we have developed the bioinformatics, data visualization software tool, Omic-Sig, which can stratify prominent phospho-substrates and their associated kinases based on differential abundances between case and control samples (e.g., tumors and their normal adjacent tissues from a cancer cohort) in a multi-omics fashion. Omic-Sig is available at https://github.com/hzhangjhu/Omic-Sig.

## Introduction

Protein phosphorylation is one of the more prominent post-translational modifications, serving an important role in regulating multiple cellular processes, such as cell growth, cell differentiation, and metabolism.^1,2^ Phosphorylation occurs when protein kinases phosphorylate their selected substrates, leading to a cascade of signaling events. Multiple reports have indicated that aberrant phosphorylation-based signaling can drive various diseases, including cancer.^1,3^

Mass spectrometry (MS) is a wide-spread analytical tool that be applied to comprehensively investigated the phosphoproteome. By conducting comparative analysis using phospho-data of paired samples (e.g., tumors and their normal adjacent tissues (NAT) in a cancer cohort), a deeper understanding of disease can be gained and lead to the identification of potential therapeutic targets.^4^ Furthermore, coupling phosphoprotemics with global proteomic and transcriptomic profiles can provide more insight into the altered expression profiles of kinases and their respective substrates.^3^

In this study, we have developed a software tool, Omic-Sig, to explore and visualize the kinase-substrate interactions by computing and ranking the differential abundances between case and control samples. Omic-Sig is constructed using C# (Microsoft Visual Studio 2017) and R (version 3.5) to provide a user-friendly interface and comprehensive data visualization.

## Methods

### Workflow of Omic-Sig

As shown in Figure 1, Omic-Sig can accept MS-based global proteomic and phosphoproteomic data as well as transcriptomic data generated from RNA-Seq as input. For each uploaded omic data set, the abundances between case and control samples (e.g., Tumor/NAT pair) are computed. By importing information on kinases and substrates, Omic-Sig will stratify kinase-substrate interactions by ranking the cases based on the calculated fold changes among different substrates of a kinase to obtain the highest ranked phosphorylation events in the majority of cases. Only phosphorylation events have more than 60% (user-adjustable) of cases with > 1.5 fold change (user-adjustable) are considered for the stratification.

**Figure 1.**
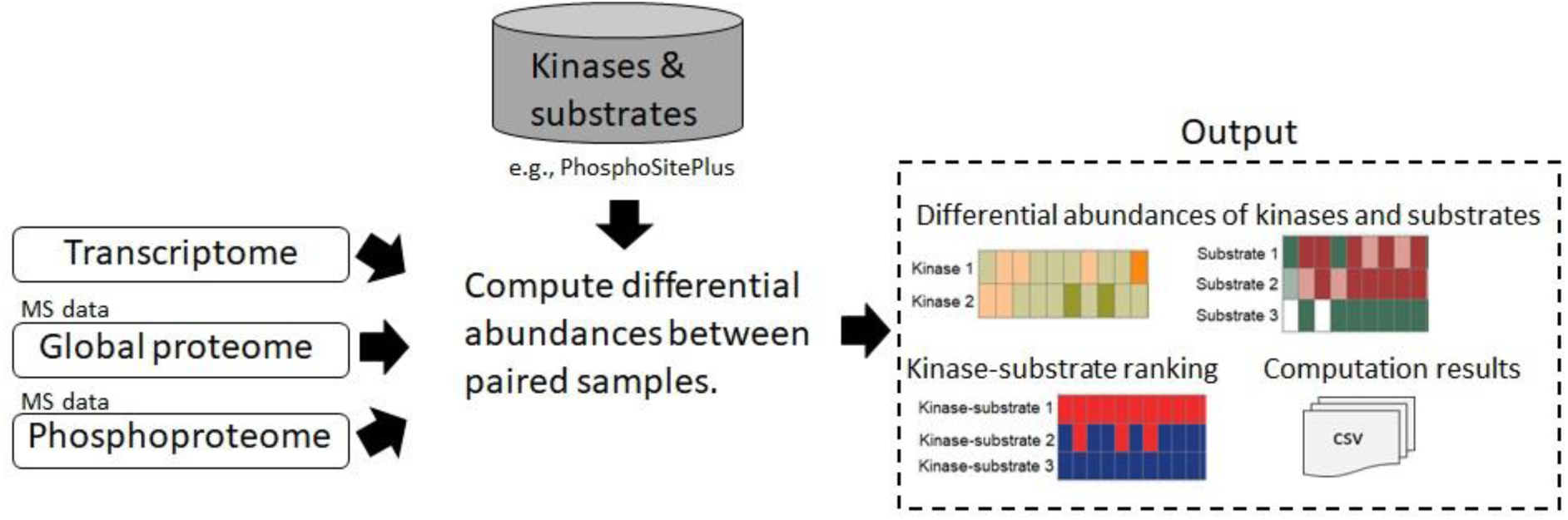
Workflow of Omic-Sig.

### Input Data

Omic-Sig requires expression matrices of global proteins and phosphosites as well as information on kinase-substrate interactions (e.g., PhosphoSitePlus^5^). It is optional to upload an mRNA expression matrix. For the expression matrices, the columns represent case or control samples, whereas each row may represent a protein, gene, or phosphosite, depending on the type of the omic data. To correctly infer the sample pairs, users need to provide a sample information table, where each row represents a sample and columns show the corresponding sample type (e.g., Tumor or NAT) and pair tag (i.e., a pair of samples is assigned the same identification tag). The expression matrix should be normalized and log-transformed prior to uploading into Omic-Sig.

### Output

We utilized a CPTAC (Clinical Proteomic Tumor Analysis Consortium) colon cancer dataset,^4^ downloaded from LinkedOmics,^6^ to exemplify the Omic-Sig pipeline. Following Omic-Sig analysis, a table of stratified kinase-substrate interactions is displayed (Figure 2) on the user-interface. Separate tables showing differential abundances between paired samples from each omic data for kinases and substrates are displayed as well (data not shown). All the computational results are automatically exported as csv files in user-specified output directory. Users can visualize the results by selecting the kinases-substrate interactions of interest (Figure 3).

**Figure 2.**
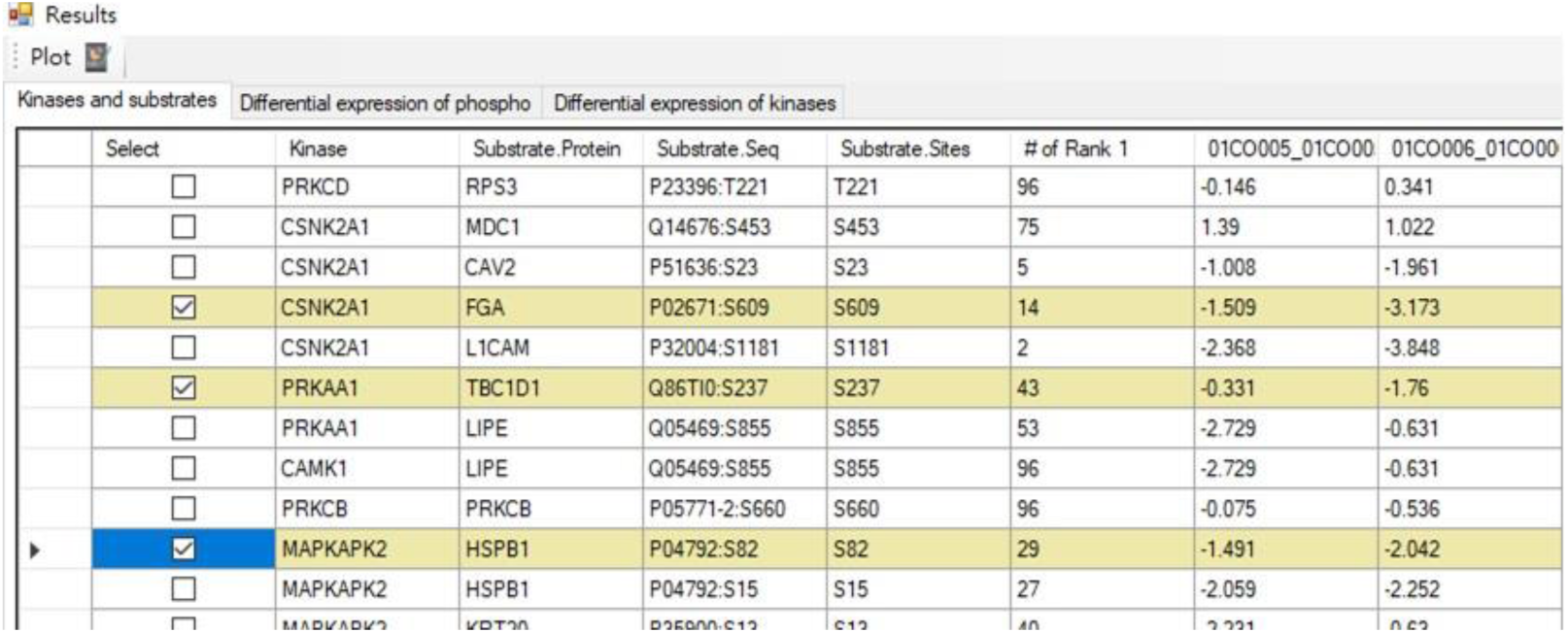
Kinase-substrate table shows kinases and their respective substrates as well as differential abundances between case and control samples for the substrates. Highlighted rows are the selected kinase-substrate interactions for visualization.

**Figure 3.**
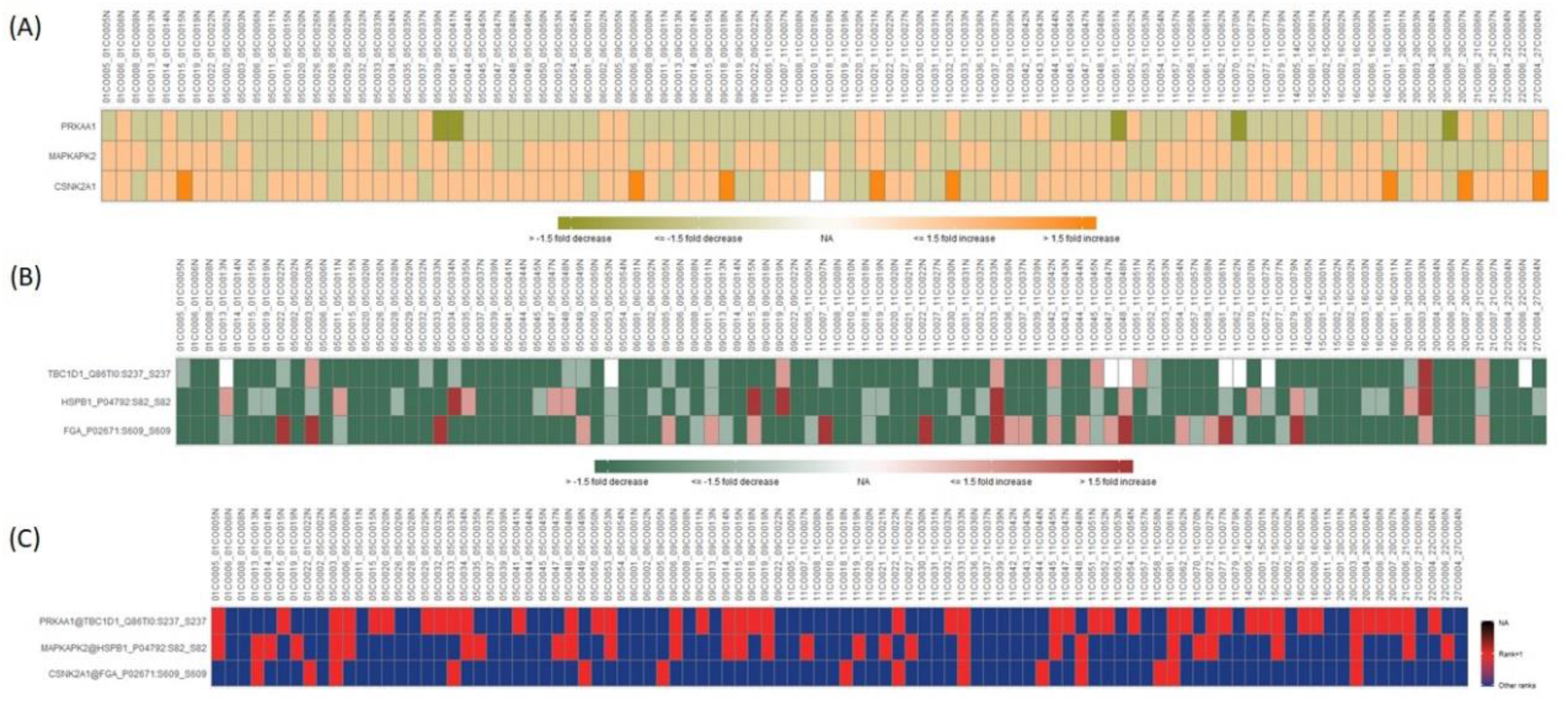
Visualizing kinase-substrate interactions. Omic-Sig generates the plots for selected kinase-substrate pairs. (A) and (B) shows differential abundances across case and control samples for selected kinases and phospho-substrates. (C) Ranked kinase-substrate events.

## Conclusions

In summary, Omic-Sig is a bioinformatics tool to stratify phospho-substrates and their associated kinases by utilizing the differential abundances between case and control samples in phosphoproteomics, global proteomics, and transcriptomics data. The tool provides data visualization on the kinase-substrate interactions allowing users to easily examine the similarity and differences across samples and different omics data. To the best of our knowledge, Omic-Sig is the first tool using paired samples to study and visualize kinase-substrate relations in multi-omics fashion.

## Funding

This work was supported by the National Institutes of Health, National Cancer Institute, the Clinical Proteomic Tumor Analysis Consortium (CPTAC, U24CA210985) and the Early Detection Research Network (EDRN, U01CA152813).

## Conflicts of Interest

The authors declare no conflict of interest.

